# An age- and time-specific space-for-time mark-recapture model

**DOI:** 10.64898/2026.07.31.742094

**Authors:** Ryan Vosbigian, Marika Dobos, Matthew R. Falcy

**Affiliations:** Department of Fish and Wildlife Sciences, University of Idaho, 875 Perimeter Dr. MS 1141, Moscow, Idaho 83844, USA; Idaho Department of Fish and Game, 3316 16th St., Lewiston, Idaho 83707, USA; U.S. Geological Survey, Idaho Cooperative Fish and Wildlife Research Unit, University of Idaho, 875 Perimeter Dr. MS 1141, Moscow, Idaho 83844, USA

**Keywords:** capture-recapture, Cormack-Jolly-Seber, Stan, migration

## Abstract

Space-for-time Cormack-Jolly-Seber (CJS) models have been developed to estimate survival of migrating animals that are imperfectly detected at spatially discrete sampling stations. However, space-for-time CJS models typically ignore survival over time and age-specific probability of movement, missing the diversity of life history strategies. To address these limitations, we extended space-for-time Cormack-Jolly-Seber models to explicitly incorporate time, age, and individual-level covariates to account for diverse life history strategies. We incorporate a sub-model that includes uncertainty in individuals’ age using a finite-mixture model. The detection and transition probabilities were parameterized using generalized linear models, facilitating flexible model specification and inclusion of spatiotemporal and individual covariates. We apply the model to detections of juvenile steelhead (*Oncorhynchus mykiss*) from two populations in the Snake River Basin in Idaho, USA, to estimate trends in survival with respect to population, age, time, and the length of individuals. Additionally, we generalized the model so that it can be applied to other systems and populations with different life history strategies and monitoring infrastructure.

## Introduction

Mark-recapture techniques are used to study population dynamics of animals. Individuals are uniquely marked and then are observed or recaptured at later occasions. Critically, non-detections can arise either from imperfect detection of individuals that are available or because the individual is unavailable because it has died or emigrated. Mark-recapture methods are commonly used to estimate animal abundance (Otis et al. 1978), survival (Cormack 1964; Jolly 1965; Seber 1965), and recruitment (Jolly 1965; Seber 1965) by modeling detectability of individuals. Multi-state mark-recapture (MSMR) models (Lebreton and Pradel 2002) are generalizations of classic Cormack-Jolly-Seber (CJS) mark-recapture models useful for incorporating (i) movement between sites (Schofield and Barker 2008), (ii) life history variation among individuals (Bonner and Schwarz 2006), and (iii) more flexible observation processes (Barker 1997; Perry et al. 2018). A specific type of MSMR model useful for migratory animals are space-for-time MSMR model (Burnham et al. 1987; Skalski et al. 1998; Buchanan et al 2015; Hance et al. 2020), where unidirectional movement past a series of fixed detection sites is substituted for successive sampling through time.

Although survival is often modeled as a discrete process over space or time, survival is a continuous process influenced by spatiotemporal conditions and individual characteristics. Time is ignored in the simple space-for-time MSMR, but time still transpires so there is need to accommodate time-varying parameters such as survival and age-dependent transitions (i.e. the animal’s probability of migrating versus remaining in place). Thus, recent advances in space-for-time MSMR models re-incorporate time into the MSMR models (Buchanan et al. 2015; Perry et al. 2018; Hance et al. 2020, 2024; Van Vleet et al. 2024). However, animal migration behavior can be complex, where movement can vary by age and individual condition (Owens et al. 2025).

For instance, salmon and steelhead (*Oncorhynchus* spp.) can move past observation sites and migrate directly or holdover (i.e. overwinter) before migrating out to the ocean, varying this behavior by age and location (Dobos et al. 2020; Owens et al. 2025). An existing model that incorporates age-specific transitions and holdovers is limited because it ignores uncertainty in age-class of individuals and is not parameterized to incorporate covariates (Buchanan et al. 2015), which limits the model’s applicability to migrating wild salmon and steelhead. Based on how wild salmon and steelhead are sampled, age-class is not directly measured for most individuals (Dobos et al. 2023). Incorporating uncertainty in individual age-class and incorporating covariates in a MSMR model would address these gaps.

We extend the framework developed by Hance et al. (2020) by developing an age- and time-specific space-for-time MSMR model. We use the observed data likelihood parameterization, which is highly computationally efficient in comparison to the complete data likelihood (Hance et al. 2020). We parameterized the state transitions and detection probabilities using generalized linear models (GLM), which allow for spatial, temporal, and individual covariates to be incorporated. We implemented the model in likelihood and Bayesian frameworks with program R (R Core Team 2025) and using Stan (Carpenter et al. 2017) to implement the Bayesian model. Our model is especially useful for species that have variable life history strategies or where movement and survival vary by location or age. We applied this model to anadromous *O. mykiss*, steelhead, (that can move directly toward the ocean or holdover) between observation sites before migrating out to the ocean. We incorporated uncertainty in individual age-class assignment by integrating the likelihood of observations over the age-assignment probability.

## Methods

We present an age- and time-specific space-for-time mark-recapture model. We made it flexible to incorporate a variety of site arrangements, site-specific covariates, temporal covariates, and some individual covariates. A constraint on site arrangement is that from each site, individuals can only transition to the next adjacent site, and backward movement is prohibited. Transitions between sites are both age- and time-specific. We parameterize the state transitions and detection probabilities using GLMs to allow for flexibility in model specification and inclusion of covariates. We start by explaining how the data are organized and then describe the various parts of the joint likelihood.

### Data description

The observed data are the individual detection histories and auxiliary information. The detection history for each individual is summarized into two matrices for instances of individuals released and recaptured (**M**) and instances of individuals released but never recaptured (**L**). Each row in the **M** matrix is a unique observation (*o*) and consists of the release site (*j_o_*), recapture site (*k_o_*) release time (*s_o_*), recapture time (*t_o_*), initial release site (*r_o_*), group for individual covariates (*g_o_*), and individual identifier (*i_o_*):

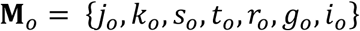

The **L** matrix consists of the final release station (*j_i_*), release time (*s_i_*), initial release station (*r_i_*), group for individual covariates (*g_i_*), and individual identifier (*i*):

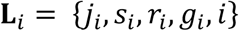

Some individuals are captured at a station but not released (i.e. removed and released elsewhere):

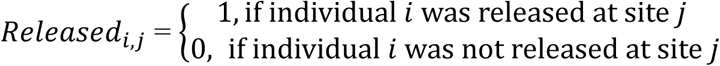

The auxiliary data include the observed age class (*ObsAge_i_*), the time stratum when age class was observed (*ObsTime_i_*), and the matrix of auxiliary data for each individual (*Z_i_*), where columns are variables and rows are for each individual.

Individuals are assumed to transition to only one site *k* from each site *j* (i.e. no forks where some individuals could transition to a different site first). For some sites *j*, individuals can delay movements before transition to the next site, whereas for other sites *j* individuals do not delay movements. This could result from the behavior of individuals or from the spatial proximity of sites. Age-classes are also constrained based on site, so that individuals can only exist at a site within a specified range of age-classes (Figure 1). The site configuration is described by *NextSite_j_* and *PrevSite_r_*_,*k*_: *NextSite_j_* = the site *k* that follows site *j*.

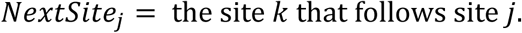

**Figure 1.**
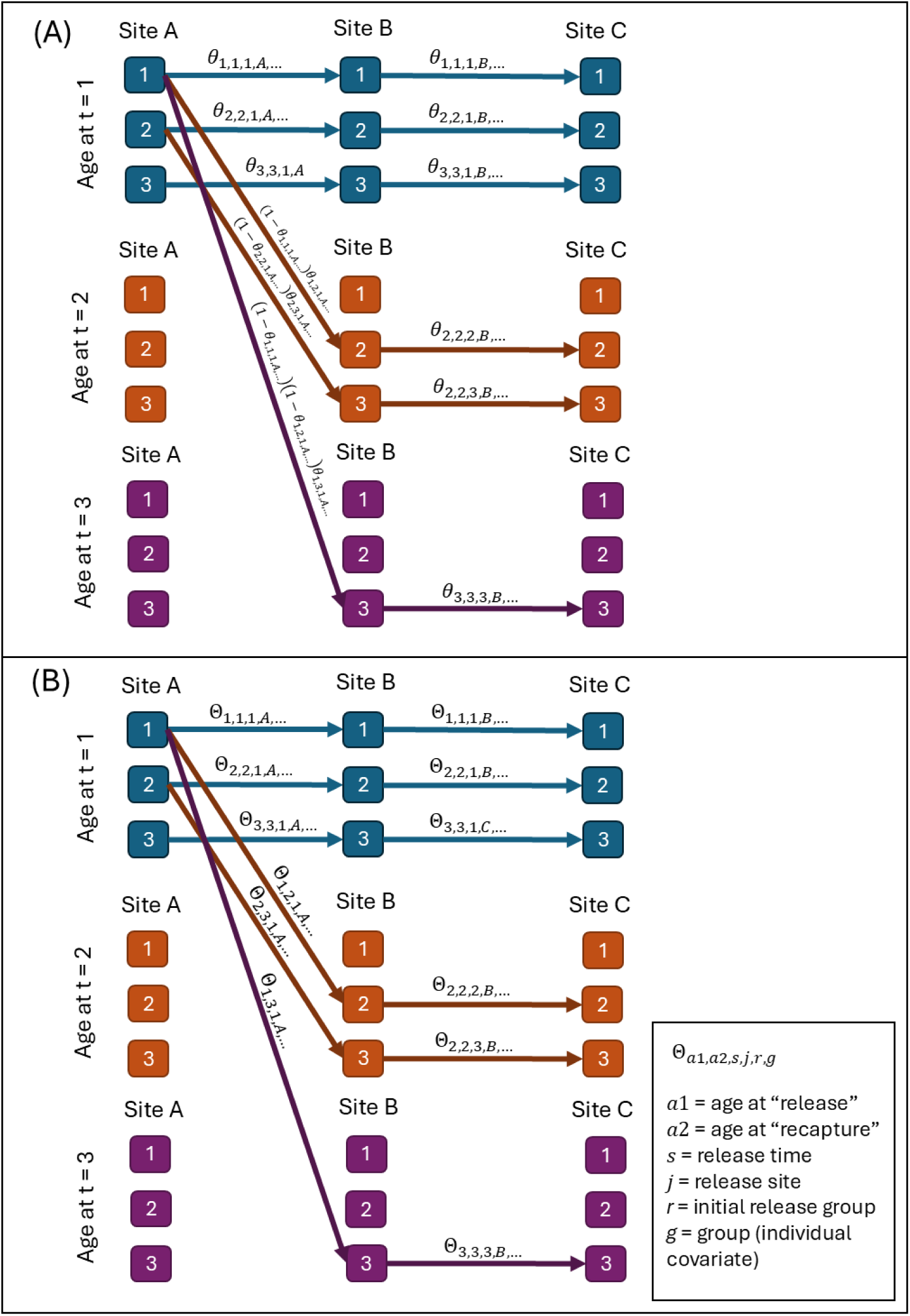
In this example design, there are three sites, individuals can be ages 1-3 years at each site, and individuals can holdover up to two years between sites A and B. The lines and parameters show the possible transitions for individuals released from site A in the first time period. The probabilities associated with the transitions are given by the conditional transition probabilities (panel A) and transition probabilities (B), where the transition probabilities are a function of the conditional transition probabilities. Individuals that do not transition are presumed to be mortalities, permanent residents, or emigrants.

*PrevSite_r_*_,*k*_ = for individuals in initial release group *r*, the site *j* that is before site *k*. Individuals from multiple initial release groups (*r*) can pass by the same site *k*, so the indices in *PrevSite_r_*_,*k*_ identify what site *j* precedes site *k* for individuals from initial release group *r*. Pairs of consecutive sites *j* and *k* where individuals can delay movements (i.e. holdover) are described by the matrix *H_j_*_,*k*_:

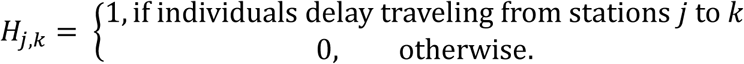

The specified minimum and maximum age classes that can exist in each site *j* are:

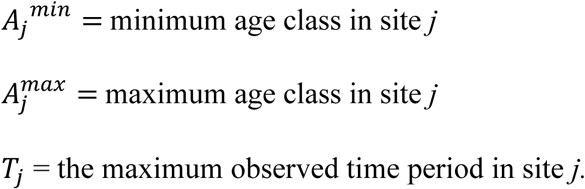

### Mark-recapture likelihood setup

The MSMR likelihood for the transition (Θ*_a_*_1,a2,*s*,*j*,*r*,*g*_) and detection (*p_a_*_1,*a*2,*t*,*k*,*r*,*g*_) probabilities is calculated using intermediary parameters. The transition probabilities, which include survival and movement probabilities, are defined as a function of conditional transition probabilities (*θ_a_*_1,a2,*s*,*j*,*r*,*g*_), which are the probability that individuals transition in a particular time interval given that they did not transition in the previous time interval (Figure 1). The indices for age (*a1* and *a2*) indicate the age (*a1*) of individuals when passing (observed or unobserved) by site *j* at time period *s* (i.e. release) and the age (*a2*) of individuals when passing by site *k* at time period *t* (i.e. recapture). The two remaining indices are for initial release site (*r*) and group (*g*). In the equation below, some indices are omitted for readability. The transition probability for individuals that move directly (in the same time interval) from one site to the next is

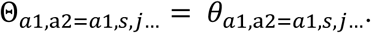

Next, transition probabilities are calculated wherein individuals holdover for one or more time intervals before passing the next site. For sites where individuals can holdover before transitioning to the next site (specified by *H_j_*_,*k*_ = 1), the transition rates are calculated from the intermediate parameters as the probability that they did not transition at a previous time. For sites where individuals only transition directly (in the same time interval) to the next site (*H_j_*_,*k*_ = 0), these transition probabilities are zero. Thus,

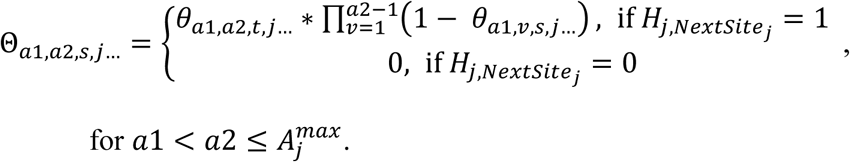

The probability that individuals never transition to the next site, apparent mortality (which includes mortality or residuals), is the product of the complement of the conditional transition probabilities 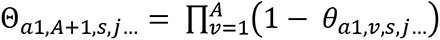.

Individuals cannot transition to a younger age, so the transition probability is zero

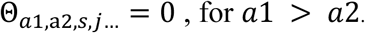

The transitions for age-classes that are lower than the minimum age-class or greater than the maximum age for a site are zero as,

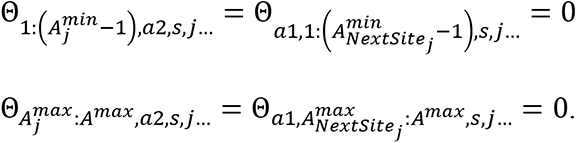

Transitions between sites that do not have holdovers are restricted to only cases where a1 and a2 are equal. If 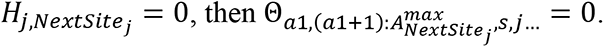

For each set of releases, which includes age, site, release time, initial release group, and individual group, the sum of the transition probabilities is defined to be 1

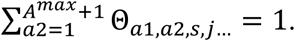

The conditional transition probabilities (*θ_a_*_1,*a*2,*s*,*j*,*r*,*g*_) and detection probabilities (*p_a_*_1,*a*2,*t*,*k*,*r*,*g*_) are parameterized using GLMs, which is described in detail in the parameterization section below. As in other CJS models, the detection probability and apparent survival for the last occasion are non-identifiable. Thus, we fixed the detection probability for the last sites to 1:

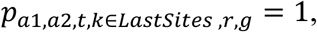

where *LastSites* is a vector of the last sites in the configuration. Non-sensible detection probabilities are set to zero:

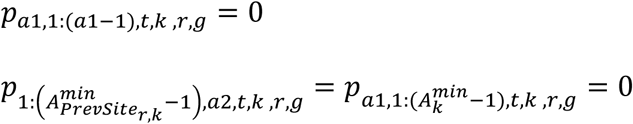

Otherwise, the values of *p_a_*_1,*a*2,*t*,*k*,*r*,*g*_ are determined by a GLM function of the data Y and parameters *ρ*:

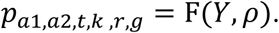

The likelihood of each pair of release and recapture is defined as *λ_j_*_,*k*,*s*,*t*,*r*,*g*,*a*1_, which is calculated from parameters recursively as in Hance et al. (2020). To obtain these values, we start with the likelihood of observations between successive capture occasions, which are given by the Θ values:

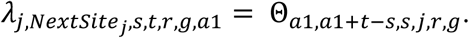

The remainder of the *λ_j_*_,*k*,*s*,*t*,*r*,*g*,*a*1_ values are the likelihood of observations between non-consecutive sites, where individuals were observed but not at the next site. The likelihood of being recaptured but not at the next site is the sum of all the possible combinations of transitions

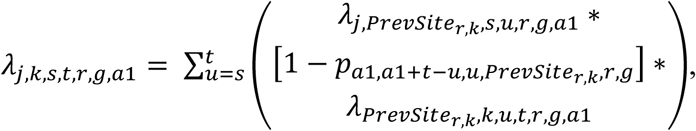

for each pair of sites *j* and *k* that are on the same path but are not consecutive (i.e. individuals that pass by site *j* pass by at least one other site before passing by site *k*). The first term in the product (*λ_j_*_,*PrevSite*_*_r_*_,*k*,*s*,*u*,*r*,*g*,*a*1_) is the likelihood of individuals transitioning from site j to the site immediately before site *k* (*PrevSite_r_*_,*k*_). The second term ([1 − *p_a_*_1,*a*1+*t*−*u*,*u*,*PrevSite*_*_r_*_,*k*,*r*,*g*_]) is the likelihood that individuals were not observed at the site immediately before site *k*. The third term (*λ_PrevSiter_*_,*k*,*k*,*u*,*t*,*g*,*r*,*a*1_) is the likelihood of individuals transitioning from the site immediately before site *k* to site *k*. The product of these three terms is computed for each combination of timing for the two transitions (from site *j* to *PrevSite_r_*_,*k*_ and from site *PrevSite_r_*_,*k*_ to site *k*) for the first transition occurring from time *s* to time *t*.

The likelihood of each release without a recapture is defined as *χ_j_*_,*s*,*r*,*g*,*a*1_, which is also calculated recursively based on intermediary parameters as in Hance et al. (2020). No individuals released at a final station (vector *LastStation*) are observed again, so

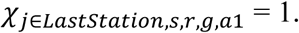

The likelihood of individuals not being observed again is the likelihood of an apparent mortality plus the sum of all the possible states where individuals survive to the next site but are not observed,

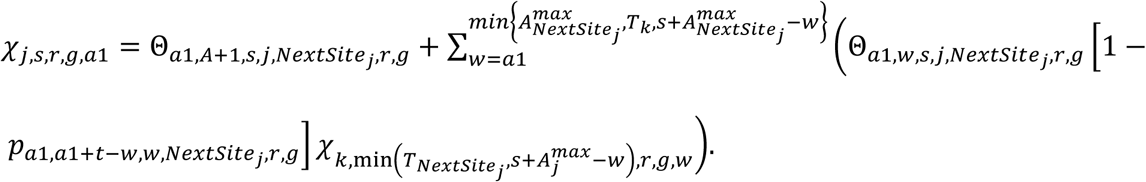

These are calculated recursively from the final stations to the first stations. The term Θ*_a_*_1,*A*+1,*s*,*j*,*NextSite*_*_j_*_,*r*,*g*_ is the probability that individuals died. In the summation, the first term Θ*_a_*_1,*w*,*s*,*j*,*NextSite*_*_j_*_,*r*,*g*_is the likelihood of individuals surviving from site *j* to the next site. The second term 1 − *p_a_*_1,*a*1+*t*−*w*,*w*,*NextSite*_*_j_*_,*r*,*g*_ is the likelihood that individuals were not detected at the site following site *j*. The third term 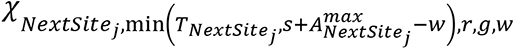 is the likelihood of individuals not being observed again after passing the site following site *j*. The products of these three terms are summed for each of the different ages that individuals could be at the site following site *j* (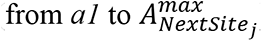). The term min (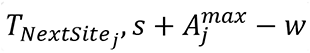) gives the maximum age that individuals can be, which incorporates the maximum time observed and the maximum age of individuals. Likewise, 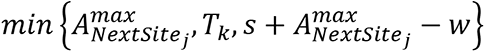 the maximum age individuals can be based on the maximum observed time period.

### Ageing uncertainty

The joint likelihood includes individuals with known ages and with unknown ages. For individuals with unknown ages, auxiliary information is used to estimate the probability of the individuals belonging to each age class. These probabilities are truncated by the maximum age class that individuals can be at specific stations.

The probability that an individual belongs to the age in a given time 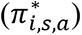 is

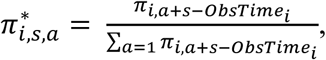

where *π_i_*_,*a*+*s*−*ObsTime*_*_i_* is the vector of age-class probabilities of individual *i* at time *s* given that it was initially observed at time *ObsTime_i_*. Alternatively, if auxiliary data for age-class were observed at a different time, then *ObsTime_i_* is when these data were collected. If the age of an individual is known, then

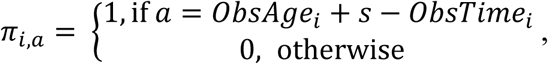

where *ObsTime_i_* is when the age-class of individual *i* was observed.

### Joint MSMR likelihood

The joint likelihood of the MSMR portion of the model is:

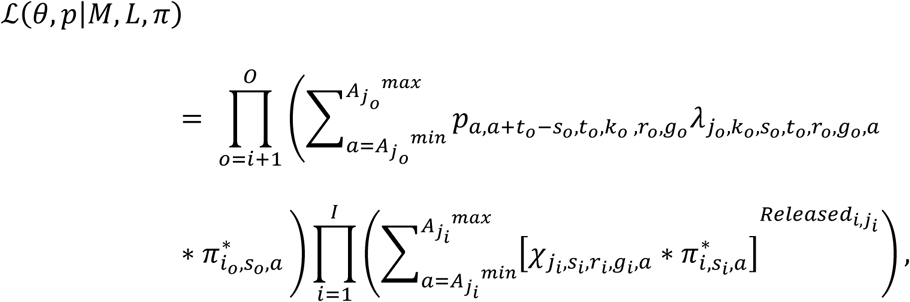

which is the product of likelihoods of each pair of release-recapture event (*p* ∗ *λ*) times the product of the likelihoods of each release with no recapture event (*χ*) integrated with respect to age-class assignment probability of individuals (*π*). Instances where individuals that were captured at an occasion but not released (where *Released_i_*_,*j*_*_i_* = 0) have a probability of no recapture of 1. The incorporation of age-class probabilities is effectively a mixture model (McLachlan et al. 2019), where the age-class probabilities are informed both by auxiliary data and detection history.

Observations with the same indices (i.e. detection history, age class covariates, and holdover duration) were grouped using an m-array (Pradel et al. 2007) to increase sampling efficiency:

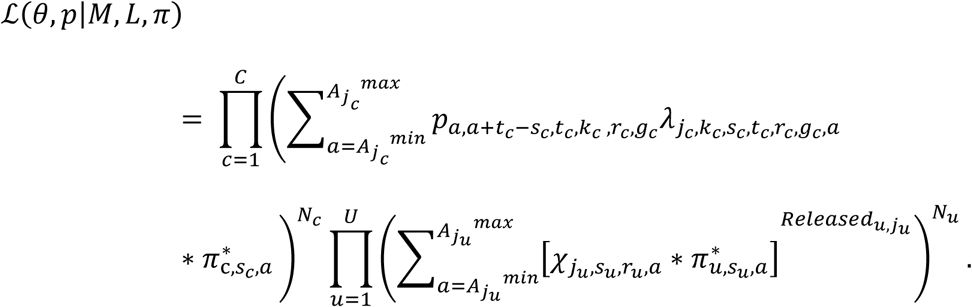

The observations of release and recaptures with the same indices (release time, recapture time, release site, recapture site, initial release site, age at release if known, age at recapture if known, age class covariates if age is unknown, and group) are indexed by *c*, and the observations of releases without recoveries (*i*) with the same indices (release time, release site, initial release site, age at release if known, age class covariates if age is unknown, and group). The number of observations grouped together is represented by *N_c_* for release and recaptures and *N_u_*for releases without recaptures, so the joint likelihood is the likelihood of the observations raised to the power of the number of observations.

### Parameterization of transition and detection probabilities

An abbreviated description of the parameterization of *θ_a_*_1,*a*2,*s*,*j*,*r*,*g*_ and *p_a_*_1,*a*2,*t*,*k*,*r*,*g*_ is given before a more detailed description. In brief, *θ_a_*_1,*a*2,*s*,*j*,*r*,*g*_ and *p_a_*_1,*a*2,*t*,*k*,*r*,*g*_ are parameterized using generalized linear models (GLM) with the logit link, where the indices *a*1, *a*2, *s*, *t*, *j*, *r*, and *g* are treated as categorical covariates and transformed into a design matrix (as in Chambers and Hastie 1992). As in GLMs, the design matrices are multiplied by a vector of **β** parameters and logit transformed to obtain the *θ_a_*_1,*a*2,*s*,*j*,*r*,*g*_ and *p_a_*_1,*a*2,*t*,*k*,*r*,*g*_values. Covariates can be included by substituting the covariate values for the indices in the design matrix. Using this parameterization, the full model, where *θ* values are unique for each conditional transition, can be expressed using notation from Chambers and Hastie (1992) as

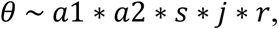

whereas a reduced model where the conditional transitions are the same for each initial release group (*r*) is

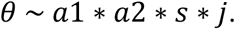

Other kinds of reduced models can be formulated using covariates. For instance, the transitions for specific sites can be fixed to be equal by including a categorical covariate with the same value for the sites. Continuous covariates can also be specified for only specific sites. Individual covariates can be included via the grouping index (*g*).

More specifically, values are parameterized by flattening the six-dimensional *θ_a_*_1,*a*2,*s*,*j*,*r*,*g*_ and *p_a_*_1,*a*2,*t*,*k*,*r*,*g*_ arrays to two-dimensional arrays with seven columns. Six of the columns are the indices associated with the *θ* or *p* value, which are the seventh column of their respective arrays. The indices are used to construct design matrices. The *θ* or *p* values are the product of the design matrix and vectors of parameters constructed using the formula on the six indices

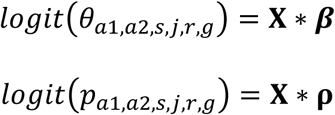

For instance, for the formula

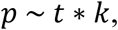

given two sites (*k* ∈ {2, 3}) and three time periods (*t* ∈ {1, 2, 3}) is equivalent to

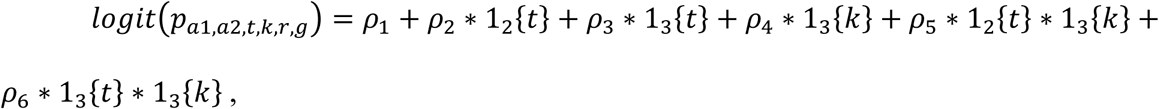

where 1*_a_*{*b*} is the indicator function:

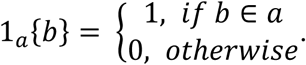

The priors for the parameters are weak and centered on zero:

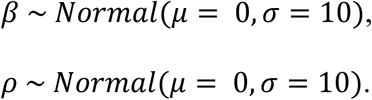

### Age class sub-model

The sub-model for age-class is an ordinal logistic regression model (Tutz 2021), which is a GLM for the probability that individuals are in observed age-classes. We used the proportional odds parameterization, which uses a logit link. The probability that the age of an individual is equal to or greater than *a* is *γ_i_*_,*a*_

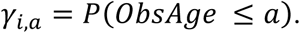

The cumulative probability (*γ_i_*_,*a*_) is estimated as

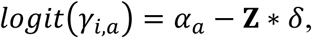

where *γ_i_*_,*a*_ is a function of thresholds between age-classes on the logit scale (*α_a_*) and linear combination of data (**Z**) and parameters (*δ*), where **Z** is a model matrix. The probability that an individual belongs to an age-class (*π_i_*_,*a*_) and the cumulative probability (*γ_i_*_,*a*_) are related were,

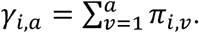

The data (Z) are a design matrix, which has a row for each individual and excludes a column for the intercept. The design matrix allows for different covariates to be included such as individual characteristics and information such as observation year or location.

The threshold parameters (*α*) are constrained so that

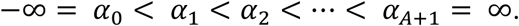

We used flat (improper) priors for the *α* parameters. For the slope (*δ*) parameters, we used weak priors centered on zero

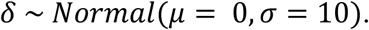

The joint likelihood for the observed age-class of individuals is according to the categorical distribution, which is the product of the age assignment probabilities for each observed individual

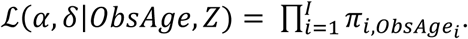

### Assumptions

The assumptions include the standard assumptions for CJS and MSMR models (Cormack 1964; Pradel et al. 2007). These include that (1) tags are not lost or misidentified, (2) capture and transitions of individuals are independent of other individuals, (3) capture and recapture does not influence transition probabilities, and (4) individuals initially captured are representative of the population. In addition to the standard CJS assumptions, other assumptions depend on the parameterization of the transition and detection probabilities. In general, all individuals from a particular initial release site (r), group (g), and age (*a1*) during the same time period (*s*) have the same transition probabilities from site *j* to *k* in time period *t*. Additionally, we assume that recapture probability is not affected by past recaptures, unless individuals were removed during sampling.

### Implementation and model evaluation

We implemented the model in a Bayesian framework and have made it publicly available as the space4time package on GitHub (Vosbigian 2026). We provide methods to calculate model fit metrics and compute derived parameters for apparent survival and transition probabilities.

Goodness-of-fit can be evaluated using a variety of different methods. We could conduct posterior predictive checks (i.e. Hance et al. 2020) where data (i.e. the **M** and **L** matrices) would be simulated from the posterior and compared to the actual observed data (Kery and Royle 2016). A Bayesian P-value could be computed from comparison of the simulated to actual data or the fit could be inspected visually (Kery and Royle 2016). However, there are many levels in the data that would make evaluating goodness-of-fit using posterior predictive checks difficult.

Alternatively, goodness-of-fit could be evaluated using the optimal goodness-of-fit tests developed by Pradel and colleagues (2007). All assumptions evaluated in Pradel et al. (2007) except one-that past detection history does not influence future detections- can be accounted for by fitting a full model for detection and transition probabilities, where the detection and state transition probability depends on the site, time period, age, holdover duration, and initial release site. Thus, goodness-of-fit can also be evaluated by fitting a full model and comparing it to reduced models using predictive performance metrics such as widely applicable information criterion (WAIC; Watanabe 2013) and Pareto-smoothed importance sampling (PSIS-LOO; Vehtari et al. 2017). Here, we evaluate goodness-of-fit using the last method, where we fit the full model and reduced models for detection probability and then compare using PSIS-LOO.

To rank and compare models, we used PSIS-LOO to estimate pointwise out-of-sample prediction accuracy (Vehtari et al. 2017; Vehtari et al. 2024). Advantages of using PSIS-LOO over WAIC include that the estimate is more robust to weak priors and influential observations and that standard errors for the estimated predictive accuracy can be approximated. Additionally, PSIS-LOO can be estimated using posterior samples of the log-likelihood of each observation.

To estimate the predictive accuracy using PSIS-LOO, we used the log likelihood with respect to individual observations rather than the log-likelihood of the m-array, so that each observation was treated as distinct.

### Methods: model application

The Potlatch River in Idaho, USA, is part of an important population of anadromous steelhead (*Oncorhynchus mykiss*). To mitigate population declines, habitat restoration is conducted in the watershed and effects on the population are monitored. There are two rotary screw traps on tributaries of the Potlatch River: Big Bear Creek and the East Fork Potlatch River. These capture juveniles as they exit the tributaries and, prior to release, technicians measure and tag fish with passive integrated transponder (PIT) tags. A portion of these individuals have scales taken, so that they can be assigned ages (Dobos et al. 2023). These tagged juveniles are detected as they migrate out to the ocean past antennas located in the hydro system by-pass facilities (Skalski et al. 1998). However, many juvenile steelhead do not migrate out directly after tagging at the rotary screw trap. Some fish holdover for a year or more before passing the first detection location, Lower Granite Dam (Dobos et al. 2020). Once fish pass Lower Granite Dam, they typically migrate out directly to the ocean without further holdovers. A key metric of interest for population monitoring and the evaluation of habitat restoration effects is the survival of juvenile steelhead from the rotary screw trap to Lower Granite Dam.

We modeled observations of *O. mykiss* captured, tagged, and released from rotary screw traps in Big Bear Creek and East Fork Potlatch River from 2008 to 2022. Data were obtained from a Columbia Basin Research Data Access query developed for a mark-recapture model, Basin TribPit DART (2026). We retained observations at any of the eight dams downriver of the Potlatch River (in order of upriver to downriver: Lower Granite, Little Goose, Lower Monument, Ice Harbor, McNary, John Day, The Dalles, and Bonneville) or in the Columbia River estuary, and then identified individuals that were removed, transported downriver by barge and, then, released downstream of Bonneville Dam. To create the detection history, we treated the rotary screw traps as the initial capture sites. The detection site at Lower Granite Dam was treated as the first recapture site, and then detection sites at Little Goose Dam and downstream were treated as the second recapture site. Pooling these observations downriver of Little Goose Dam increased the sample size of detections and reduced the number of parameters required to fit the model, which helped avoid convergence issues.

To fit models, we used a sequential approach where we started with age-class only models to select the best age-class sub-model, then fit the mark-recapture model to test various detection probability formulas, and, finally, included individual covariates in the mark-recapture model. We used the space4time package to fit all the models. For the age-class only models, we evaluated five different models, fitting them using maximum likelihood and compared fit based on AIC (Table 1). Using the top age-class sub-model and the full conditional transition probability model, we fit the mark-recapture model with five different parameterizations of detection probability (Table 1).

**Table 1.**
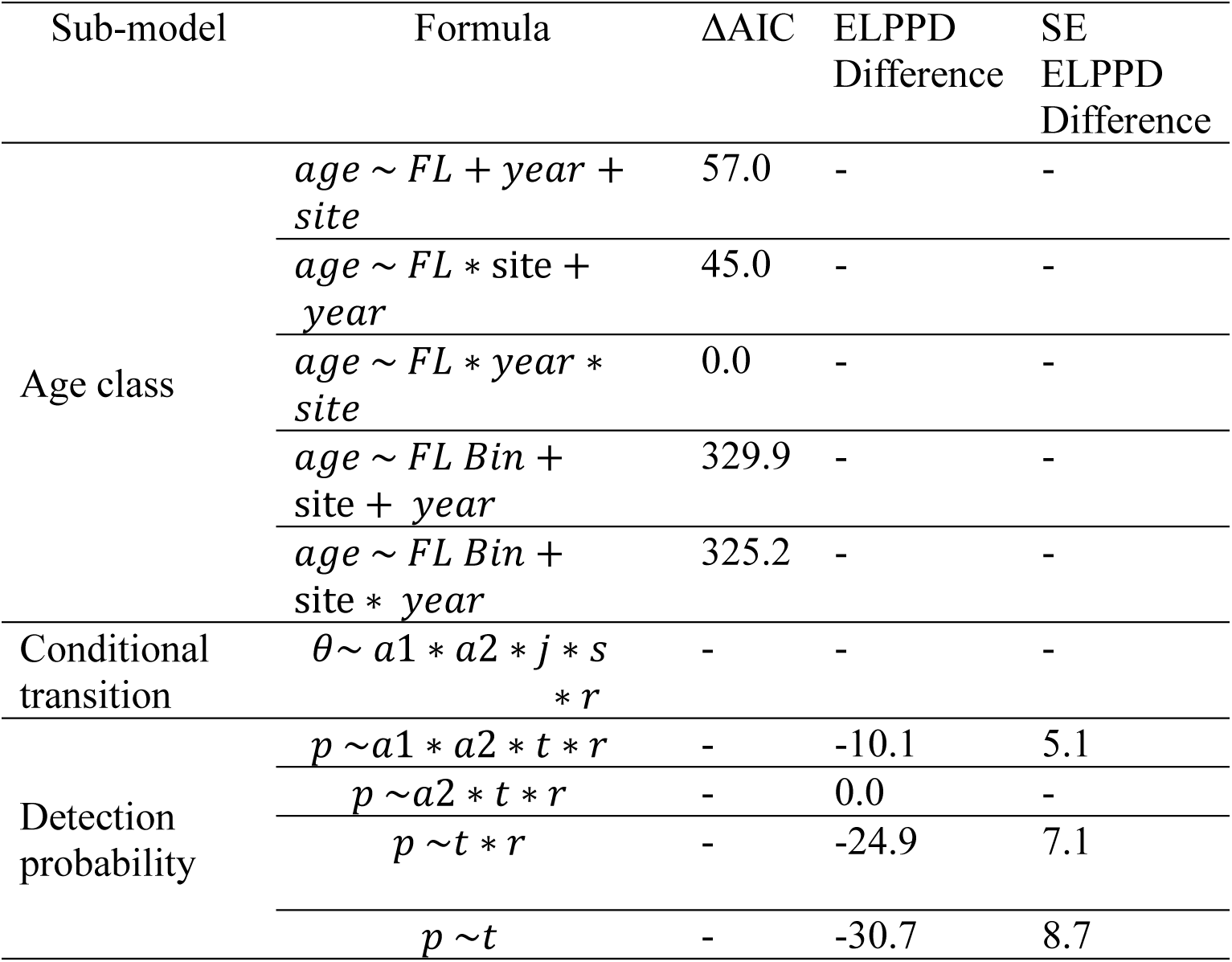
Initial model selection table. The age-class only models were fit using maximum likelihood and are compared using AIC. The full mark recapture models were fit using Bayesian methods and are compared using PSIS-LOO.

| Sub-model | Formula | ΔAIC | ELPPD<br>Difference | SE<br>ELPPD<br>Difference |
| --- | --- | --- | --- | --- |
| Age class | $age \sim FL + year + site$ | 57.0 | - | - |
| | $age \sim FL * site + year$ | 45.0 | - | - |
| | $age \sim FL * year * site$ | 0.0 | - | - |
| | $age \sim FL \text{ Bin} + site + year$ | 329.9 | - | - |
| | $age \sim FL \text{ Bin} + site * year$ | 325.2 | - | - |
| Conditional transition | $\theta \sim a1 * a2 * j * s * r$ | - | - | - |
| Detection probability | $p \sim a1 * a2 * t * r$ | - | -10.1 | 5.1 |
| | $p \sim a2 * t * r$ | - | 0.0 | - |
| | $p \sim t * r$ | - | -24.9 | 7.1 |
| | $p \sim t$ | - | -30.7 | 8.7 |

Next, we included covariates. We included a group covariate for individual lengths measured when captured at rotary screw traps. We binned lengths into 10 groups of equal sample size using quantiles. We modeled the relationship between the average length within each group and conditional transitions from the rotary screw traps to Lower Granite Dam. The effect of fork length on conditional transitions was included as an interaction between age when released from the rotary screw trap and site. Fork length was modeled in two different ways, linear and quadratic, with respect to site and age (Table 2). We fit all mark-recapture models using Stan with 3 chains with 1000 burn-in iterations and 1000 additional iterations for the posterior. We compared the fitted mark-recapture models using PSIS-LOO.

**Table 2.** Models including individual covariates for length-at-age were compared using PSIS- LOO.

| Conditional transition<br>formula | ELPPD<br>Difference | SE<br>ELPPD<br>Difference |
| --- | --- | --- |
| $\theta \sim a1 * a2 * j * s * r$ | -257.7 | 30.5 |
| $\theta \sim a1 * a2 * j * s * r$<br>+ *firsttransition*: $r * a1$<br>* *FL* | -40.3 | 9.0 |
| $\theta \sim a1 * a2 * j * s * r$<br>+ *firsttransition*: $r * a1$<br>* *FL* + *firsttransition*: $r$<br>* $a1 * FL^2$ | 0 | - |

## Results

The best model for age-class, based on AIC, included a three-way interaction between fork length, observation year, and release site. We included this sub-model for age-class in the full mark-recapture models. Using PSIS-LOO, the best model for detection probability included different detection probabilities depending on initial release site (r), “recapture” observation time (t), and age during “recapture” event (a2). This model was a substantially better fit compared to the other models by -10.1 (s.e. = 5.1) expected log pointwise predictive density (ELPPD) (Table 1). The top covariate model included fork lengths as a quadratic varying by population and age and was substantially the best model by 40.3 ELPPD (s.e. = 9.0).

To generate population-specific estimates of transition probabilities and apparent survival, we used the mark-recapture model without individual covariates. The transition probabilities and apparent survival probabilities varied by site (initial release site and recapture sites), year, and age class (Figures 2 and 3). In both populations, older fish tend to migrate directly rather than holdover. Younger fish from the East Fork Potlatch River tended to holdover a year, whereas most fish in Big Bear Creek transitioned directly downriver. From both populations, there have been declines in apparent survival over time. We found that apparent survival was generally positively related to fork length but the effect of fork length varied by population and age (Figure 4). For most ages, the effect of increasing fork length was greater at smaller sizes than larger sizes.

**Figure 2.**
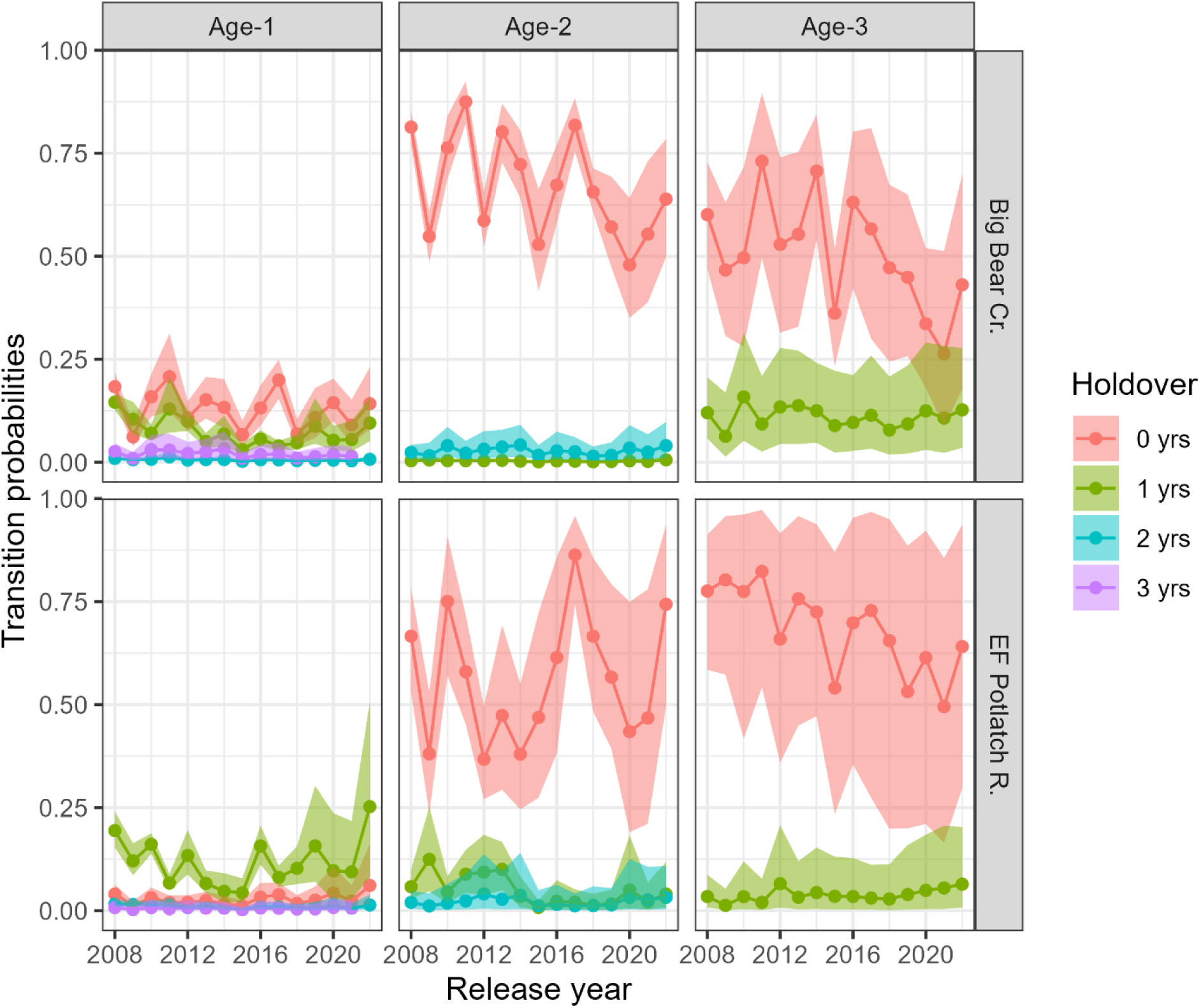
The transition probabilities and holdover duration of individuals from Big Bear Creek and the East Fork Potlatch River to Lower Granite Dam varies by the year and age they were released from the sites. These transition probabilities represent the probability that an individual traveled to Lower Granite Dam in the same year as initial capture or held-over for 1 or more years. The points are the mean transition probabilities, and the shaded regions are the 95% credible intervals.

**Figure 3.**
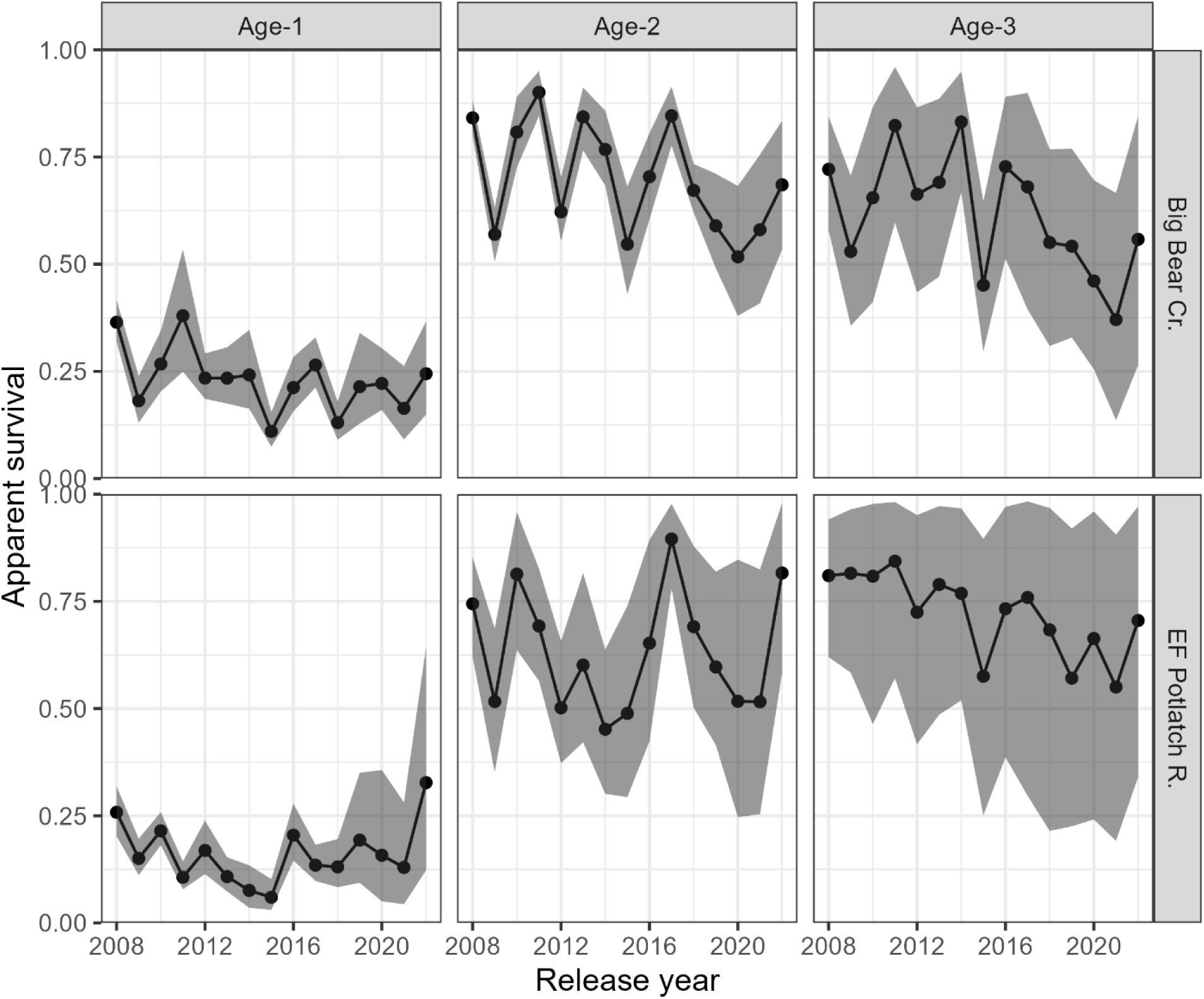
The apparent survival of individuals from Big Bear Creek and the East Fork Potlatch River to Lower Granite Dam varies by the year and age they were released from the sites. The points are the mean apparent survival, and the shaded regions are the 95% credible intervals.

**Figure 4.**
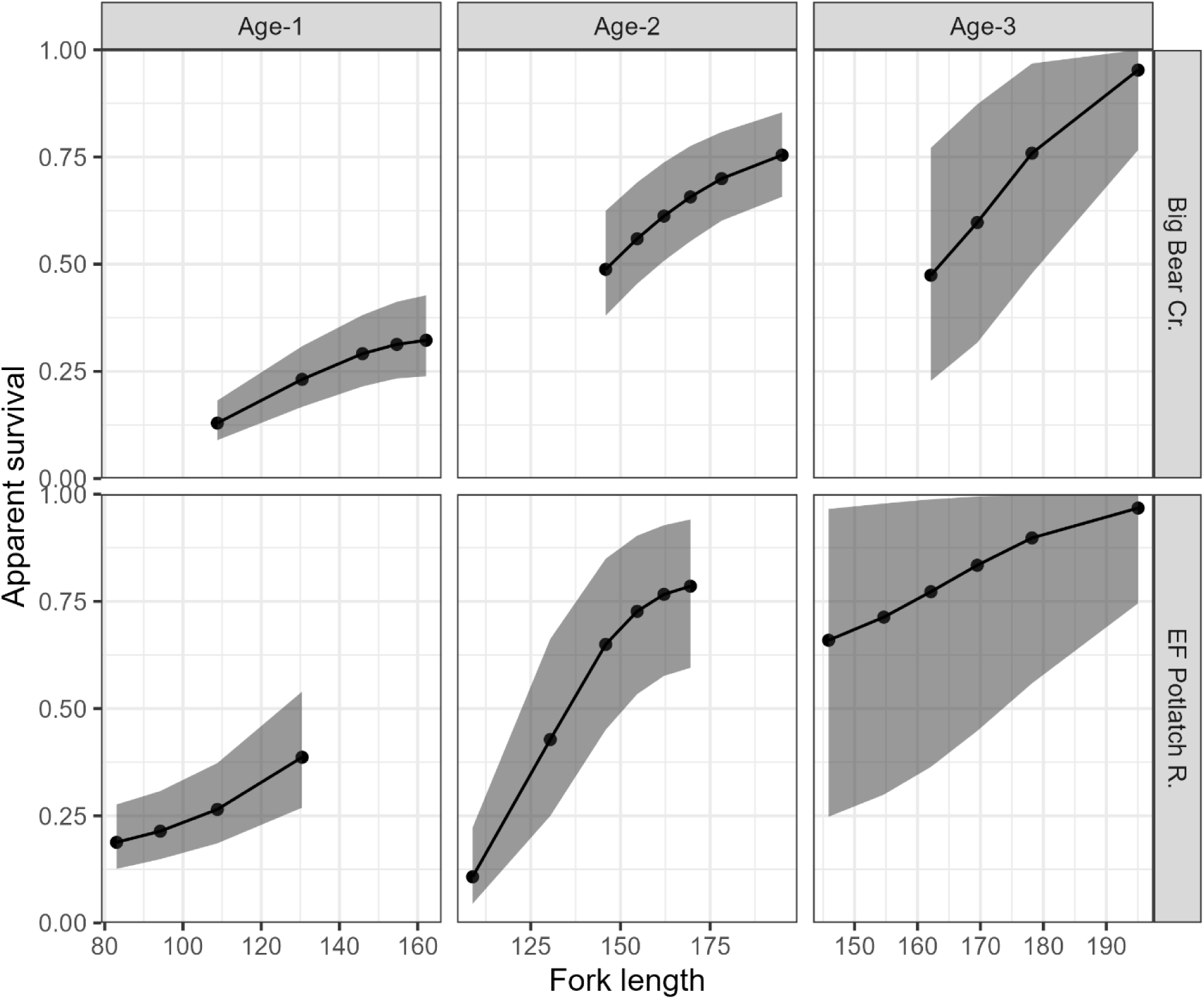
Apparent survival from Big Bear Creek and the East Fork Potlatch River to Lower Granite Dam increased as individual fork length-at-age (mm) increased. The effect shows the survival for release year 2016. The points are the mean apparent survival, and the shaded regions are the 95% credible intervals. The range of fork lengths for each age-class are the 10% and 90% quantile of fork length for each age class for each population.

Using the m-array allowed us to fit the models efficiently, by condensing the 37,560 observations to 3,961 values when evaluating the likelihood for the top model.

## Discussion

Here, we present a novel MSMR model that explicitly accounts for ageing uncertainty in an age-specific space-for-time mark-recapture model. We parameterized the likelihood in terms of the observed data likelihood by applying methods developed in Hance et al. (2020). We parameterized the detection and conditional state transition probabilities using logistic regression, allowing for many kinds of models to be fit. For instance, substituting more complex time-varying or age-specific detection probabilities for a simpler detection probability parameterization is straightforward to incorporate. Additionally, we incorporated individual covariates for individual body condition. Our implementation and inclusion of individual covariates allowed us to efficiently implement the observed data likelihood. We gained efficiency in sampling by using an m-array, where observations with identical capture histories and auxiliary data were grouped together.

The flexibility of our MSMR model allows for many kinds of site configuration and model parameterizations to be evaluated using the same framework as long as assumptions can be met. This resulted in a limited parameterization in some respects. For instance, we did not differentiate between survival and movement in the conditional transition rates, and each sub- model is only parameterized by fixed effects. Further extensions to our model could incorporate hierarchical parameters or splines (Lang and Brezger 2004; Bonner and Schwarz 2011) to the detection and state transition probabilities.

A challenge in MSMR models is parameterizing the state transitions that comprise movement and survival. Here, we parameterized the state transitions using conditional state transitions, which are the probability that an individual transitions from one site to the next given that it did not transition to that next site in the previous time periods. We did not differentiate between movement and survival. However, Hance et al. (2020) assumed that survival was constant, but arrival probabilities were dependent on time. More specifically, Hance et al. (2020) assumed that the survival of all fish that passed a station at a specific interval was the same for all fish, regardless of travel time. This may not always be reasonable, for instance, when the range of travel times is broad.

Other parameterizations that attempt to separate survival and arrival times may not be identifiable. There must be some assumptions about movement and survival to estimate them separately. Horton et al. (2011) presented a similar model for Atlantic salmon (*Salmo salar*), where they estimated true survival and movement as opposed to apparent survival. However, it is safe to assume that Atlantic salmon that survive would at some point emigrate past monitoring infrastructure. This assumption is invalid for steelhead, which are only partially migratory, with some steelhead permanently residing as freshwater residents whereas others migrate to the ocean (Kendall et al. 2015).

By explicitly accounting for ageing uncertainty in the mark-recapture likelihood, we were able to overcome important challenges. A particular challenge in estimating survival of juvenile salmonids in the Columbia River Basin is that the age of all individuals is often presumed fixed and known (Buchanan et al. 2015). In practice, the age of individuals was predicted using a multinomial model (Dobos et al. 2023), and then individuals with observations that differed from known life history strategies were dropped from the analysis. For instance, if age-3 fish from Big Bear Creek were assumed to migrate directly to Lower Granite Dam, then any individuals assigned age-3 would be removed from analysis if they were observed to holdover a year. This could result in biased estimates for older age-classes because observations were removed.

Additionally, this could bias estimates if the transition rates vary substantially over age by mixing different ages together. We included the sub-model for age and explicitly incorporated the age-specific probabilities so that we can separate out the differences between different ages. We parameterized the influence of age-class uncertainty effectively as a mixture model (McLachlan et al. 2019), where the mixing proportions are informed by the age sub-model as well as the detection history.

Our age- and time-specific MSMR model will help researchers infer drivers of survival and movement rates for emigrating salmonids while accounting for their complex life history strategies. The estimated transition probabilities indicated a diversity of migration strategies exhibited across age groups, where some ages were more likely to migrate directly (Figure 2). We found that fork length was positively related to survival across ages and that the magnitude of the effect depended on age (Figure 4). Fork length and age (because age is positively related to fork length) have generally been found to be positively related to survival of juvenile salmonids (Evans et al. 2014; Buchanan et al. 2015). Understanding the effect for fork length and length-at-age on survival is important for evaluating the effectiveness of restoration actions (Chen et al. 2023). If an action increases length-at-age of steelhead by some amount, then we can understand how much their survival can be expected to increase.

We designed a novel statistical model to help fill a gap between existing data collection infrastructure and protocols, and practical resource management needs. Future work could use this model to evaluate the effects of environmental and biotic factors on steelhead survival.

## Acknowledgments

We thank Timothy Copeland and Ryan Kinzer and for helpful suggestions on the manuscript. Any use of trade, firm, or product names is for descriptive purposes only and does not imply endorsement by the United States Government.

